# In silico analysis of Missense Variants in Human MPL Gene Associated with Essential Thrombocythemia

**DOI:** 10.1101/830372

**Authors:** Sahar G. Elbager, Abier A. Makkawi, Hadeel A. Mohamed, Fauzia A. Abdelrahman, Lamia H. Osman, Moroj F. Hameed, Asia M. Elrashid, Mohamed Y. Basher, Safinaz I. Khalil, Magdi Bayoumi

## Abstract

**Introduction:** The proto-oncogene (MPL) gen encodes the receptor for thrombopoietin (TPO-R), a member of hematopoietic receptor superfamily. Thrombopoietin (TPO), the primary cytokine regulating self-renewal of hematopoietic stem cells, thrombopoiesis and megakaryocytopoiesis. TPO binding to TPO-R induces activation of Janus Kinase 2 (JAK2). Activated JAK2 triggers the activation of downstream positive signaling pathways, leading to the survival, proliferation, and differentiation of hematopoietic cells. Mutations in MPL gene possibly will alter the normal regulatory mechanisms. Numerous MPL mutations have been observed in various hematopoietic cancers such as essential thrombocythemia and primary myelofibrosis and leukemias. In this study, we performed a comprehensive in silico analysis of the functional and structural impact of non-synonymous (nsSNP) that are deleterious to TPO-R structure and function.

**Methodology:** The data on human MPL gene was retrieved from dbSNP/NCBI. Nine prediction algorithms; SIFT, Polyphen, PROVEAN, SNAP2, Condel, PhD-SNP, I-Mutant, Mutpred. RaptorX and Chimera were used to analyzing the effect of nsSNPs on functions and structure of the TPO-R. STRING and KEGG database were used for TPO-R protein-protein interaction.

**Results and Discussion:** As per dbSNP database, the human MPL gene contained 445 missense mutations. A total 5 nsSNPs (D295G, R257C, Y252H, R537W and D128Y) were predicted to have the most damaging effects on TPO-R structure and function. STRING and KEGG revealed that MPL had strong interactions with proteins that involved in cell growth, apoptosis, signal transduction pathway, some cancers pathways such as colorectal cancer, lung cancer, pancreas cancers, and skin cancer. A literature search revealed that Y252H has contribute to the development of essential thrombocythemia.

**Conclusion:** These in silico predictions will provide useful information in selecting the target SNPs that are likely to have functional impact on the TPO-R and moreover could act as potential targets in genetic association studies. Keywords: In Silico analyses; JAK2; Missense Variants; MPL gene; Thrombopoietin (TPO); Single nucleotide polymorphism (SNP).

## Introduction

Essential thrombocythemia (ET) is one of the myeloproliferative neoplasms (MPNs), characterized by a clonal expansion of megakaryocytes, genomic instability, dysregulated signaling pathways and subsequent overproduction of inflammatory markers [1-3]. MPNs patients have the potential to transform into acute myeloid leukemia (AML) or myelodysplastic syndrome (MDS) [4,5]. The most frequent genes involved are JAK2, Calreticulin (CALR) and myeloproliferative leukemia virus oncogene (MPL) mutations of ET patients [6-8]. Here, we predict the influence of various SNPs in MPL gene use in silico tools predict the influence of missense variants in the MPL genes.

The human myeloproliferative leukemia virus proto-oncogene (MPL) gene, a cellular homologue of the oncogene v-mpl, belongs to the haematopoietic receptor superfamily, located on chromosome one (1p34.2) composed of 12 exons [9,10]. The protein encoded by MPL gene, Thrombopoietin receptor (TPO-R), cytokine receptor superfamily, is a 95 kD has an amino acid length of 635 consists of four functional domains, including putative signal peptide, extracellular domain, transmembrane domain and the intracellular domain [11, 12]. The extracellular domain (ECD) it is composed of two adjacent pairs of fibronectin-III (FNIII)-like domains known as a cytokine receptor module (CRM) forms the interface for cytokine binding [12]. The transmembrane domain (TMD) is a helical section of the receptor that anchors it to the cell surface that anchors it to the cell surface [12]. The cytoplasmic domain, likely largely unstructured, binds JAKs and signal transduction [12].

The endogenous ligand for TPO-R (thrombopoietin-Tpo) is the primary cytokine regulating megakaryocytopoiesis, thrombopoiesis and self-renewal of haematopoietic stem cells, as reflected by expression of the TPO-R on megakaryocytes, platelets and hematopoietic progenitors [13-15].

TPO-R lacks intrinsic kinase activity and utilizes the Janus kinase (JAK) protein family to transduce a signal from the extracellular cytokine to the nucleus within the cell. JAK2 associated with the cytoplasmic domain of TPO-R and get activated by phosphorylation upon TPO signaling. Consequently, the activated JAKs phosphorylate the receptor and the signal transducer itself, prompts the activation of downstream positive signaling pathways, including JAK-STAT and mitogen-activated protein kinase (MAPK) and phosphatidylinositol-3 kinase (PI3K) pathways [16], leading to the survival, proliferation, and differentiation of hematopoietic cells [17, 18].

Mutations in MPL gene could alter the normal controlling mechanisms; some may cause an independent activation of TPO-R and the development of thrombocytosis refractory anaemia with ringed sideroblasts associated with marked thrombocytosis (RARS-t) and myeloproliferative neoplasms (MPNs) [19], while other types of mutations result in a complete or partial loss of TPO-R function, thus leading to thrombocytopenia congenital amegakaryocytic thrombocytopenia (CAMT) [20].

In the present study we aimed to determine the influence of various SNPs in MPL gene using in silico prediction software and assessing the effect of these SNPs on the structure and function of TPO-R protein that may have an important role in disease susceptibility.

## 2. Material and method

### 2.1 Retrieval of SNP ids

The data on human MPL gene (Gene ID: 4352) was retrieved from the Entrez Gene databases from National Center for Biological Information (NCBI) database on 13 March 2019. The MPL protein sequence (accession ID: P40238) and SNPs information of the MPL gene were obtained from UniProtKB databases (http://www.uniprot.org) and NCBI dbSNP (http://www.ncbi.nlm.nih.gov/snp/) respectively.

### 2.2 Sequence homology-based prediction of deleterious nsSNPs by using SIF

**SIFT** Sorting intolerant from tolerant server was used to identify the tolerated and deleterious SNPs. The effect of amino acid substitution on protein structure was assessed on the basis of degree of conservation of amino acids using sequence homology Substitution of an amino acid at each position with probability < 0.05 is predicted to be deleterious and intolerant, while probability ≥ 0.05 is considered as tolerant, nsSNPs within dbSNP retrieved data were selected as an input for SIFT (http://siftdna.org/www/SIFT_dbSNP.html) [21].

### 2. 3 Prediction of Functional effect of non synonymous SNP by Provean

PROVEAN (Protein Variation Effect Analyzer; http://provean.jcvi.org/index.php) is used to predict the possible impact of a substituted amino acid and indels on protein structure and biological function. It analyses the nsSNPs as deleterious or natural, if the final score was below the threshold score of −2.5 were considered deleterious; scores above this threshold were considered neutral [22]. The input query is a protein FASTA sequence along with amino acid substitutions

### 2. 4 Functional significance of substitution by Polyphen2

Polyphen -2 (polymorphism and phenotype; http://genetics.bwh.harvard.edu/pph2/) server was used to predict the functional impact of amino acid substitution on protein structure and function based on sequence based characterization. The Prediction outcome was obtained in the form of probability score which classifies the variations as ‘probably damaging’, ‘possibly damaging’ and ‘benign’ [23]. UniProt protein accession ID P40238 along with position and name of wild type and variant amino acids of screened nsSNPs were submitted as query.

### 2.5 SNAP2

(Screening of Non-Acceptable Polymorphism 2; https://rostlab.org/services/snap2web/) is a tool, developed based on a neural network classification method which is freely available. It predicts the effect of nsSNPs on protein function [24]. The input query submitted is the protein FASTA sequence and lists of mutants which provided scores of each substitution that can then be translated into binary predictions neutral or non-neutral effect natural.

### 2.6 Condel

(CONsensus DELeteriousness score of missense SNVs; http://bbglab.irbbarcelona.org/fannsdb/query/condel). Condel is a method, used assess the outcome of non-synonymous SNVs, via applying a CONsensus DELeteriousness score that combines various tools. It computes a weighted approach of missense mutations from the complementary cumulative distributions of scores of deleterious and neutral mutations [25]. The input query submitted is the Ensembl Protein Id ENST00000372470 along with the position and name of wild type and variant amino acids of screened nsSNPs.

### 2.7 Prediction of Disease Associated SNPs by PhD SNP

PhD-SNP (Predictor of human Deleterious Single Nucleotide Polymorphisms; http://snps.biofold.org/phd-snp/phd-snp.html) web server was used to predict if a given single point protein mutation can be classified as a disease-related or as neutral polymorphism. This server was mainly based on the support vector machines which an corroborate all the information regarding variations from the existing databases [26]. The input FASTA sequences of protein along with the residues change were submitted to PhD-SNP server for the analysis.

### 2.8 Prediction of nsSNPs Impact on the Protein Stability by I-Mutant2.0

I-Mutant2.0 is a tool used for prediction of changes in protein stability due to single site mutations under different conditions (http://gpcr.biocomp.unibo.it/cgi/predictors/I-Mutant2.0/I-Mutant2.0.cgi). It is a web server based on support vector machine which worked on dataset derived from Protherm, a database of experimental records on protein mutations. It can predict the stability changes in protein with 80% accuracy based on its structure and with 77% of accuracy based on its sequence [27]. The input can be submitted as either in the form of protein sequence or on a structure basis. For the present study, input was submitted in the form of protein FASTA sequence.

### 2.9 Identification of Functional SNPs in Conserved Regions by ConSurf

Conserve (https://consurf.tau.ac.il) web-server calculates the evolutionary conservation of amino acid substitution in proteins [28]. Consurf gives the output in the form of score where score 9 represent the most conserved and 1 represent the highly variable amino acid.

### 2.10 Analyzing the Effect of nsSNPs on Physiochemical Properties by Mutpred

Mutpred was used to predict structural and functional changes as a consequence of amino acid substitution. (http://mutpred.mutdb.org/). These changes were expressed as probabilities of gain or loss of structure and function. In addition, it predicts molecular cause of disease. The MutPred output contains a general score (g), i.e., the probability that the AAS is deleterious/disease-associated and top five property scores (p), where p is the P-value that certain structural and functional properties are impacted. A missense mutation with a MutPred (g) score > 0.5 could be considered as “harmful,” while a (g) score > 0.75 should be considered a high confidence “harmful” prediction [29]. The input was submitted in the form of protein FASTA sequence along with amino acid substitution

### 2.11 Prediction of structural effect of point mutation on the protein sequence using Project HOPE

HOPE, Have Our Protein Explained (HOPE; http://www.cmbi.ru.nl/hope/home) is an easy-to-use web service that analyzes the structural effects of a point mutation in a protein sequence. FASTA sequence of whole protein and selection of mutant variants were submitted to project hope server. HOPE server predicts the output in the form of structural variation between mutant and wild type residues [30].

### 2.12 Homology modelling using RaptorX and UCSF Chimera

RaptorX (http://raptorx.uchicago.edu/StructurePrediction/predict/), a protein structure prediction server, predicts 3D structures for protein sequences lacking close homologs in the Protein Data Bank (PDB). RaptorX uses a non-linear scoring function to combine homologous information with structural information for a given protein Sequence [31]. FASTA protein sequence was submitted to RaptorX server to get the model as PDB file. The resultant PDB files was opened using Chimera program which was used to visualize the PDB structure. UCSF Chimera (http://www.cgl.ucsf.edu/chimera). UCSF Chimera is a highly extensible program for interactive visualization and analysis of molecular structures and related data, and hence modifies the original amino acid with the mutated one to see the impact that can be produced [32]. The outcome is then a graphic model depicting the mutation.

### 2.11 Prediction of Genetic and Protein Interactions by STRING and KEGG

Protein-protein interactions are important to assess all functional interactions among cell proteins. STRING (Search Tool for the Retrieval of Interacting Genes/Proteins; https://string-db.org/) [33]. The STRING database gives a protein-protein interaction either it is direct or indirect associations. The input option we use is the protein name and the organism. KEGG (Kyoto Encyclopedia of Genes and Genomes; http://www.genome.jp/kegg/) [34] is a knowledge base for systematic analysis of gene functions in terms of the networks of genes and molecules, including metabolic pathways, regulatory pathways, and molecular complexes for biological systems.

## 3. Results

### 3.1. SNP Dataset from dbSNP

As per dbSNP database, the human MPL gene investigated in this work contained a total of 3587SNPs: 346 SNPs in 3′ UTR region, 166 SNPs in 5′ UTR region, 2439 SNPs in intron region, 623 SNPs in coding synonymous regions, 6 SNPs splice acceptor, 6 SNPs splice donor, 53 SNPs non coding transcript variant, one terminator codon variant,, 24 stop gained mutations and 445missense mutations. The missense nsSNPs were selected for the investigation.

### 3.2. Prediction of Functional Mutations

A total number of 445 nsSNPs MPL gene were submitted as batch to the SIFT program. According to our SIFT analysis predicted, 41 SNPs were predicted to be deleterious, 32 SNPs were predicted to be tolerated and 371 nsSNPs were not found.

Our PROVEAN analysis predicted that 7 nsSNPs were deleterious and 34 nsSNPs were neutral. Polyphen-2 results analysis predicted that 24 nsSNPs were probably damaging, whereas 8 nsSNPs were predicted to be ‘possibly damaging’ and the remaining 9 nsSNPs was categorized as benign. The SNAP2 Analyzer predicted that 22 nsSNPs were affected the function of protein and 19 nsSNPs were neutral. CONDEL analysis showed that 31 SNPs were predicted to be deleterious, 10 SNPs were predicted to be neutral. PhD-SNP predicted 23 nsSNPs were neutral and 18 nsSNP to be associated with disease (Table 1).

**Table 1:** List of nsSNP analysis by SIFT, PolyPhen-2, PROVEAN, SNAP2, CONDEL, PHD-SNP.

| NID | Residue | SIFT | PROV | POLY-2 | SNAP2 | CONDEL | PHD-SNP |
| --- | --- | --- | --- | --- | --- | --- | --- |
| 6088 | E168K | Deleterious | Neutral | Probably Damaging | Neutral | Deleterious | Disease |
| 3820551 | R537Q | Deleterious | Neutral | Probably Damaging | Effect | Deleterious | Disease |
| 28928907 | R102P | Deleterious | Neutral | Probably Damaging | Effect | Neutral | Disease |
| 28928908 | P275T | Deleterious | Deleterious | Probably Damaging | Effect | Deleterious | Neutral |
| 61754776 | P70L | Deleterious | Neutral | Possibly Damaging | Neutral | Neutral | Neutral |
| **113696793** | **D295G** | **Deleterious** | **Deleterious** | **Probably Damaging** | **Effect** | **Deleterious** | **Disease** |
| 117656396 | L265F | Deleterious | Neutral | Benign | Neutral | Neutral | Neutral |
| **121913611** | **R257C** | **Deleterious** | **Deleterious** | **Probably Damaging** | **Effect** | **Deleterious** | **Disease** |
| 121913612 | P635L | Deleterious | Deleterious | Probably Damaging | Effect | Deleterious | Neutral |
| 137952228 | T183M | Deleterious | Neutral | Probably Damaging | Neutral | Deleterious | Neutral |
| 138450235 | G614A | Deleterious | Neutral | Probably Damaging | Effect | Deleterious | Neutral |
| 139770726 | E335K | Deleterious | Neutral | Probably Damaging | Effect | Neutral | Neutral |
| **141311765** | **Y252H** | **Deleterious** | **Deleterious** | **Probably Damaging** | **Effect** | **Deleterious** | **Disease** |
| 142161843 | S557A | Deleterious | Neutral | Benign | Neutral | Neutral | Neutral |
| 144210383 | G238C | Deleterious | Neutral | Probably Damaging | Effect | Deleterious | Neutral |
| 145714475 | F45S | Deleterious | Neutral | Probably Damaging | Effect | Deleterious | Disease |
| 146120160 | G467S | Deleterious | Neutral | Probably Damaging | Neutral | Deleterious | Disease |
| 147608148 | S402P | Deleterious | Neutral | Benign | Neutral | Deleterious | Disease |
| 147851301 | L580S | Deleterious | Neutral | Benign | Neutral | Deleterious | Neutral |
| 148276667 | R75H | Deleterious | Neutral | Probably Damaging | Effect | Deleterious | Neutral |
| **148784027** | **R537W** | **Deleterious** | **Deleterious** | **Probably Damaging** | **Effect** | **Deleterious** | **Disease** |
| 149810307 | V368L | Deleterious | Neutral | Possibly Damaging | Neutral | Neutral | Neutral |
| 150004498 | V556F | Deleterious | Neutral | Benign | Effect | Deleterious | Neutral |
| 188798468 | P2L | Deleterious | Neutral | Probably Damaging | Neutral | Deleterious | Neutral |
| 200454070 | R426Q | Deleterious | Neutral | Probably Damaging | Effect | Deleterious | Disease |
| **201101813** | **D128Y** | **Deleterious** | **Deleterious** | **Probably Damaging** | **Effect** | **Deleterious** | **Disease** |
| 201998783 | R351C | Deleterious | Neutral | Probably Damaging | Neutral | Deleterious | Disease |
| 202137992 | P173A | Deleterious | Neutral | Possibly Damaging | Neutral | Neutral | Neutral |
| 202206935 | L109F | Deleterious | Neutral | Benign | Neutral | Neutral | Disease |
| 267598614 | E99K | Deleterious | Neutral | Possibly Damaging | Neutral | Neutral | Neutral |
| 267598615 | E149K | Deleterious | Neutral | Possibly Damaging | Effect | Deleterious | Neutral |
| 367594982 | S625A | Deleterious | Neutral | Probably Damaging | Neutral | Deleterious | Disease |
| 368753117 | R71Q | Deleterious | Neutral | Benign | Effect | Neutral | Neutral |
| 369486165 | E332K | Deleterious | Neutral | Possibly Damaging | Effect | Deleterious | Neutral |
| 369504259 | D545G | Deleterious | Neutral | Benign | Effect | Deleterious | Disease |
| 370346540 | A203T | Deleterious | Neutral | Benign | Effect | Deleterious | Neutral |
| 371227552 | P133L | Deleterious | Neutral | Possibly Damaging | Neutral | Deleterious | Neutral |
| 371232858 | A418T | Deleterious | Neutral | Probably Damaging | Neutral | Deleterious | Neutral |
| 372164360 | F315I | Deleterious | Neutral | Probably Damaging | Neutral | Deleterious | Disease |
| 373621350 | R390C | Deleterious | Neutral | Probably Damaging | Effect | Deleterious | Neutral |
| 374829965 | V538L | Deleterious | Neutral | Possibly Damaging | Neutral | Deleterious | Disease |

Because each computational method uses different parameters to evaluate the nsSNPs, concordance was done. A total of 5 substitutions (D295G, R257C, Y252H, R537W and D128Y) were found to deleterious by all the algorithms used in the study

### 3. 3 Prediction of nsSNPs Impact on the Protein Stability by I-Mutant2.0

The prediction of stability changes of selected 5 nsSNPs by I-Mutant 2.0 is given in Table 2. All SNPs were predicted to decrease effective stability of the TPO-R.

**Table 2:** Prediction result of I-Mutant software.

| SNP ID | Residue | Stability Prediction | Reliability Index (RI) | Ph | Temp |
| --- | --- | --- | --- | --- | --- |
| rs113696793 | D295G | Decrease | 3 | 7.0 | 25 |
| rs121913611 | R257C | Decrease | 6 | 7.0 | 25 |
| rs141311765 | Y252H | Decrease | 7 | 7.0 | 25 |
| rs148784027 | R537W | Decrease | 6 | 7.0 | 25 |
| 201101813 | D128Y | Decrease | 3 | 7.0 | 25 |

### 3. 4 Analysis of Conservation profile

ConSurf analysis predicted D128 to be exposed and conserved residue, D295, R257 and R537 and are exposed and highly conserved residue, i.e., a functional residue, which suggests that these positions are important for the TPO-R function. ConSurf analysis also predicted Y252 to be buried and conserved residue, i.e., a structural residue (Table 3, Fig. 1).

**Table 3:**
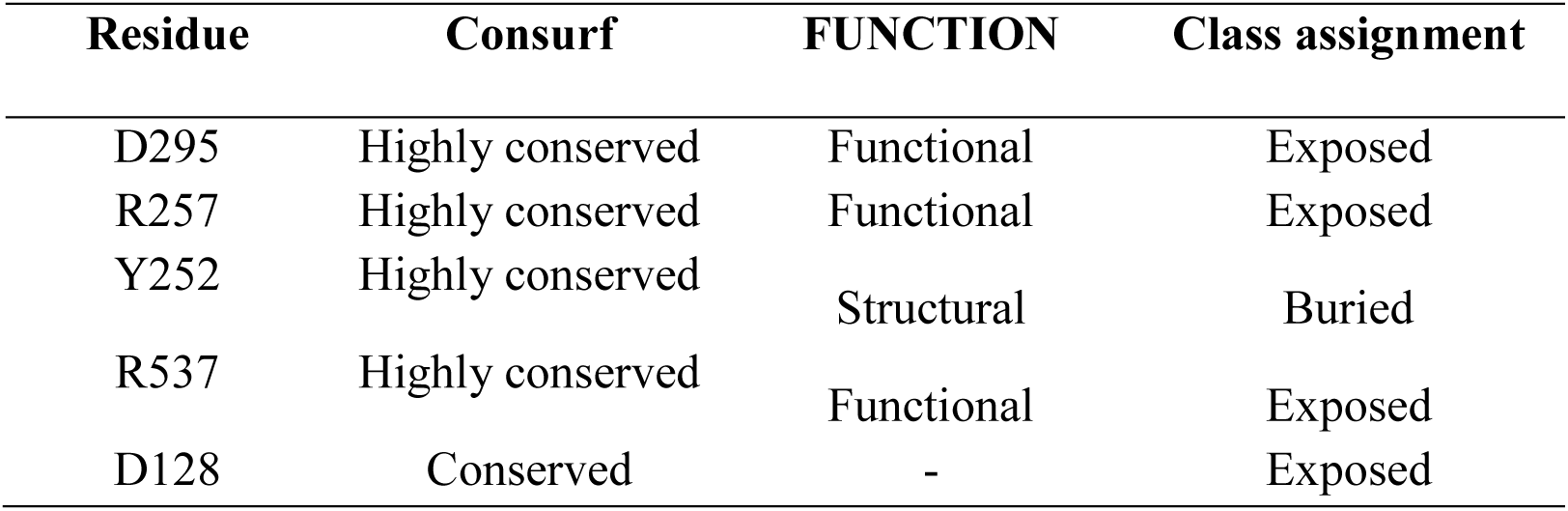
Conservation profile of amino acids in TPO-R.

**Fig. 1.**
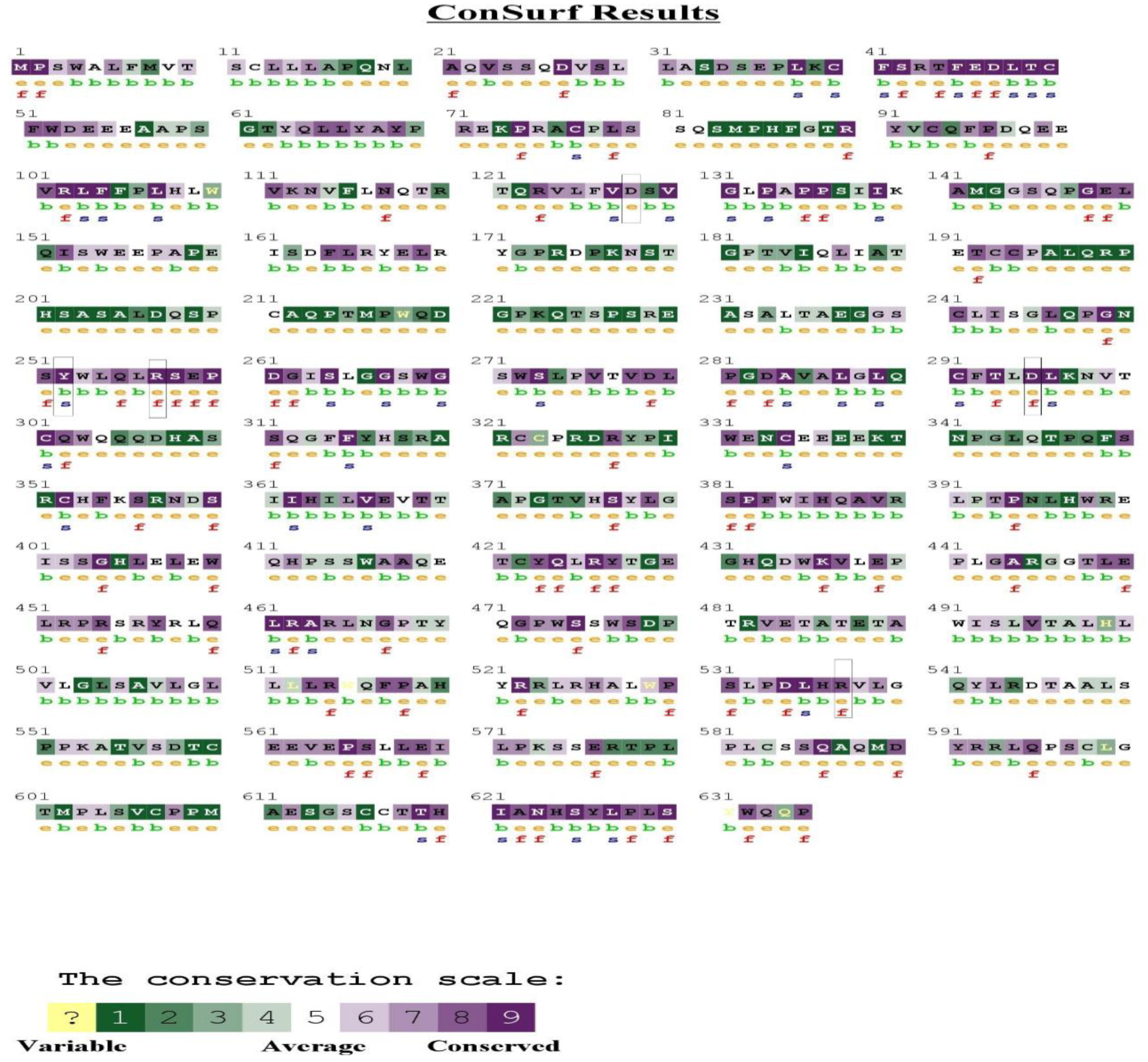
Analysis of evolutionary conserved amino acid residues of **TPO-R** by ConSurf. The color coding bar shows the coloring scheme representation of conservation score

### 3.5 Analyzing the Effect of nsSNPs on Physiochemical Properties Mutpred

MutPred result showed (R257C, Y252H and R537W) were highly harmful, D295G and D128Y were harmful. The top possible molecular mechanism disrupted were predicted; loss of sheet (P = 0.0817) for the mutation D295G, loss of disorder (P = 0.0414) for the mutation R257C, gain of disorder (P = 0.0223) for the mutation Y252H and gain of catalytic residue at R537 (P = 0.1186) for the mutation R537W, gain of phosphorylation at D128 (P = 0.054) for the mutation D128Y. Table 4 summarizes the result obtained from MutPred server.

**Table 4.**
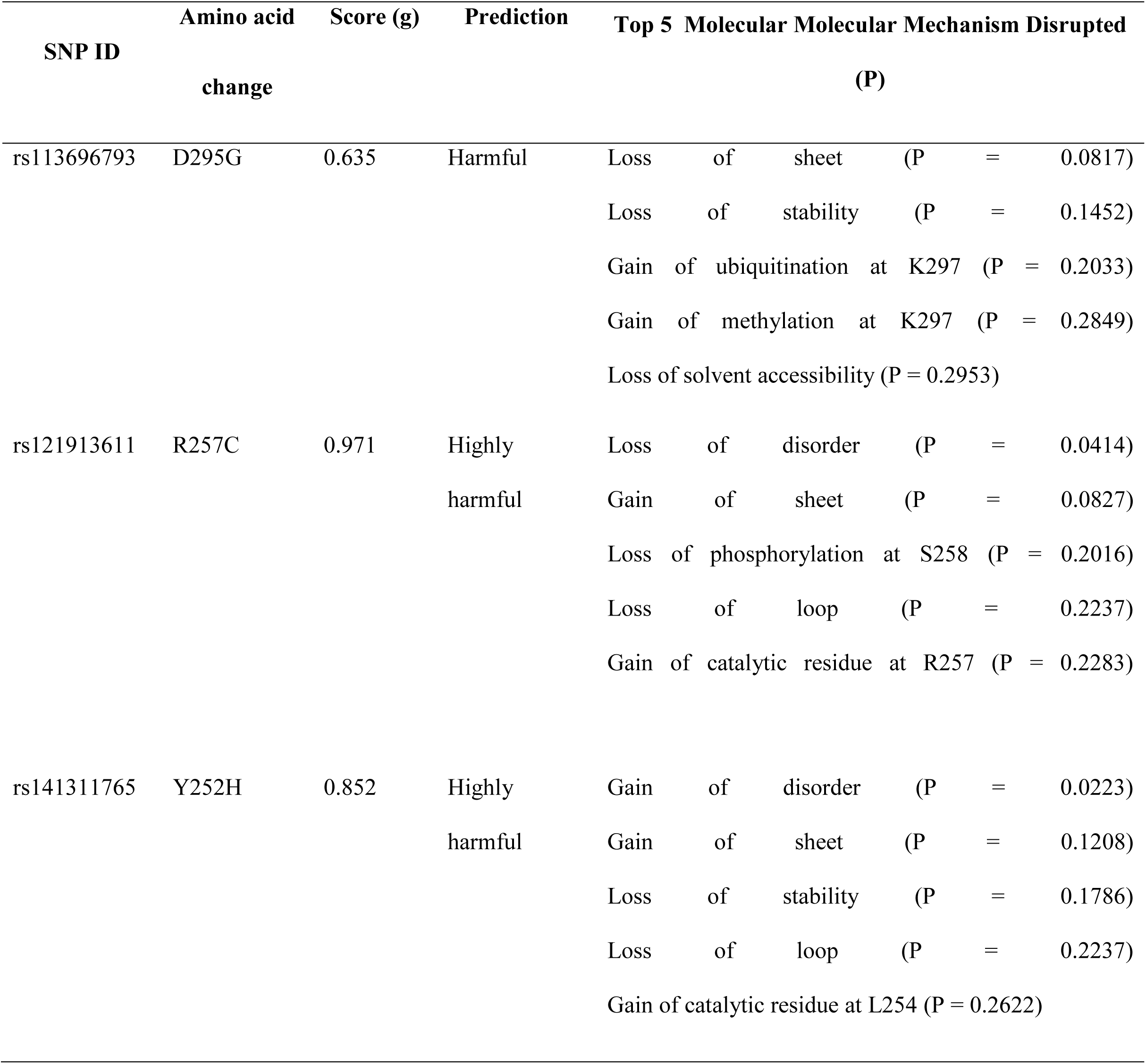
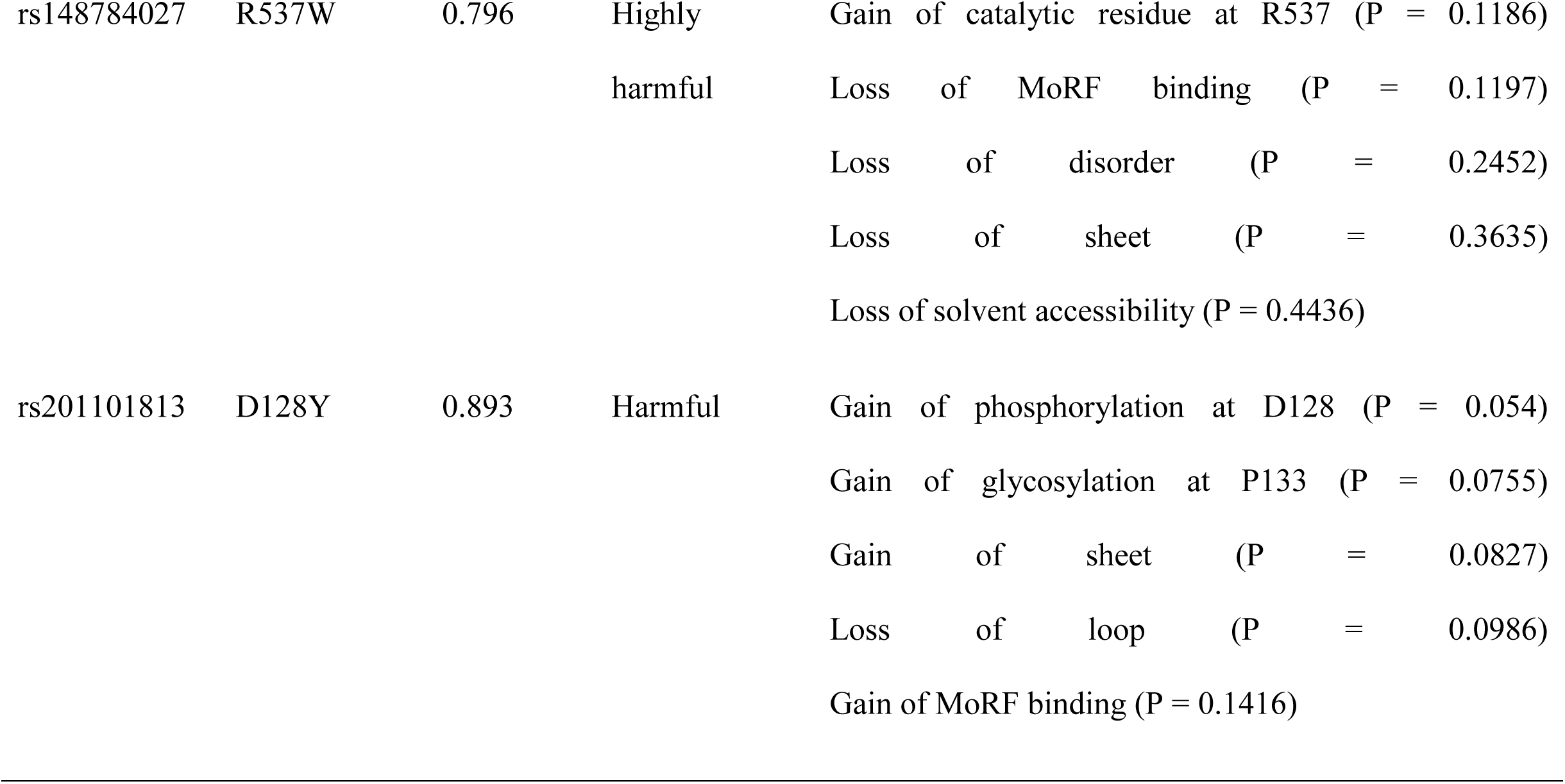
Analysis of the effect of nsSNPs in TPO-R structure, function, and evolution by MutPred server.

| SNP ID | Amino acid change | Score (g) | Prediction | Top 5 Molecular Mechanism Disrupted (P) |  |
| --- | --- | --- | --- | --- | --- |
| rs113696793 | D295G | 0.635 | Harmful | Loss of sheet | (P = 0.0817) |
|  |  |  |  | Loss of stability | (P = 0.1452) |
|  |  |  |  | Gain of ubiquitination at K297 | (P = 0.2033) |
|  |  |  |  | Gain of methylation at K297 | (P = 0.2849) |
|  |  |  |  | Loss of solvent accessibility | (P = 0.2953) |
| rs121913611 | R257C | 0.971 | Highly harmful | Loss of disorder | (P = 0.0414) |
|  |  |  |  | Gain of sheet | (P = 0.0827) |
|  |  |  |  | Loss of phosphorylation at S258 | (P = 0.2016) |
|  |  |  |  | Loss of loop | (P = 0.2237) |
|  |  |  |  | Gain of catalytic residue at R257 | (P = 0.2283) |
| rs141311765 | Y252H | 0.852 | Highly harmful | Gain of disorder | (P = 0.0223) |
|  |  |  |  | Gain of sheet | (P = 0.1208) |
|  |  |  |  | Loss of stability | (P = 0.1786) |
|  |  |  |  | Loss of loop | (P = 0.2237) |
|  |  |  |  | Gain of catalytic residue at L254 | (P = 0.2622) |

|  |  |  |  |  |
| --- | --- | --- | --- | --- |
| rs148784027 | R537W | 0.796 | Highly | Gain of catalytic residue at R537 (P = 0.1186) |
|  |  |  | harmful | Loss of MoRF binding (P = 0.1197) |
|  |  |  |  | Loss of disorder (P = 0.2452) |
|  |  |  |  | Loss of sheet (P = 0.3635) |
|  |  |  |  | Loss of solvent accessibility (P = 0.4436) |
| rs201101813 | D128Y | 0.893 | Harmful | Gain of phosphorylation at D128 (P = 0.054) |
|  |  |  |  | Gain of glycosylation at P133 (P = 0.0755) |
|  |  |  |  | Gain of sheet (P = 0.0827) |
|  |  |  |  | Loss of loop (P = 0.0986) |
|  |  |  |  | Gain of MoRF binding (P = 0.1416) |

### 3.6 Analyzing the structural effect of point mutation on the protein sequence

The project HOPE was used to investigate the structural effects of these amino acid substitutions. Its results showed that D295G, R257C, Y252H and R537W are highly conserved and their substitutions are probably damaging to the protein structure. For example, the D128Y was changes to amino acid bigger than the residue in wild type while D295G, R257C, Y252H and R537W were changes to smaller amino acid. D295G, R257C, R537W and D128Y resulted in change in the net charge of TPO-R (Table 5). It is known that protein charge and mass affect spatio-temporal dynamics of protein-protein interaction; hence, these changes could alter the ability of TPO-R to interact the other proteins and cytokines.

**Table 5.**
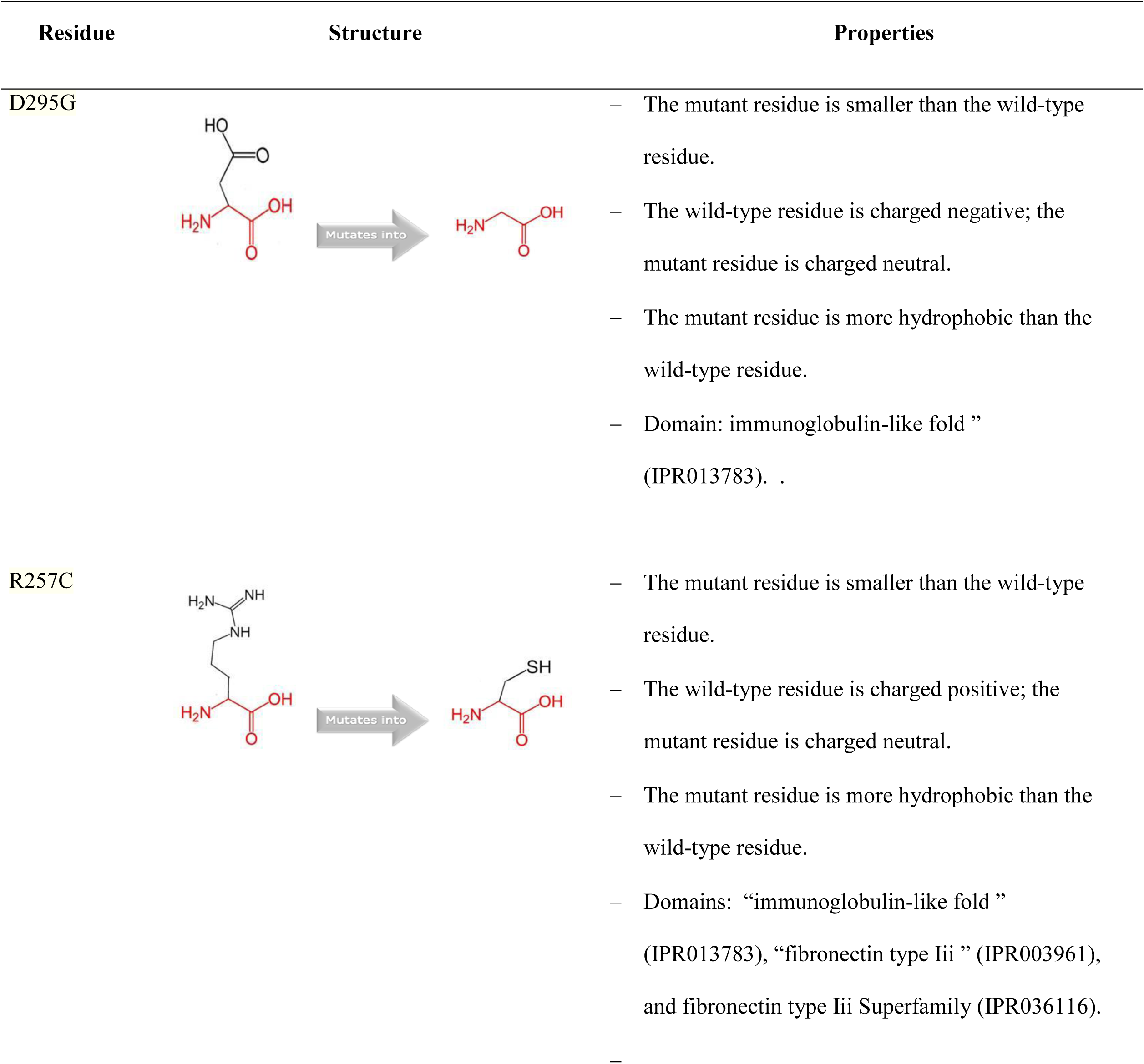
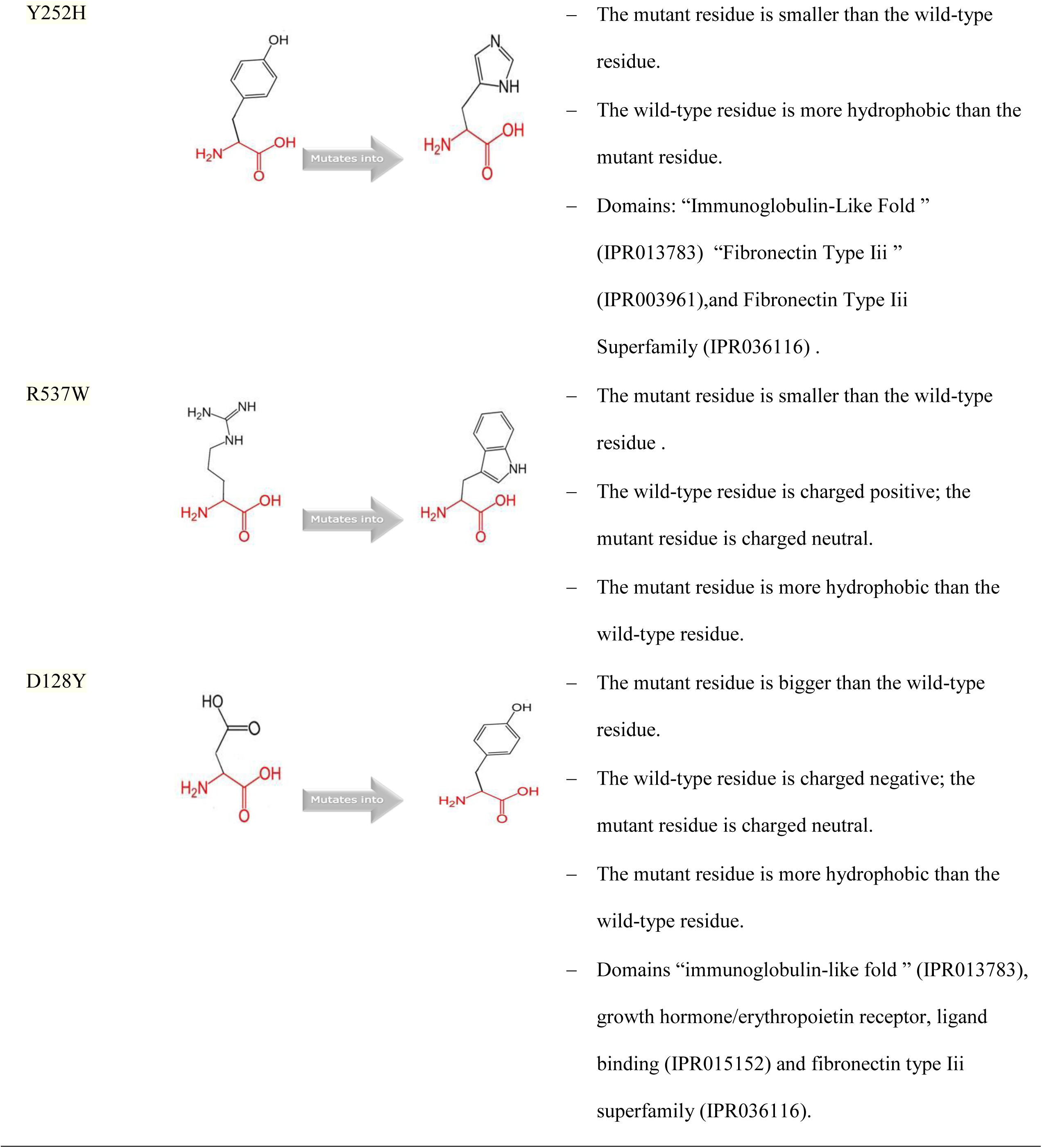
Schematic structures of the wild-type residue (left) and mutant residue (right) amino acid for each Mutation.

### 3.7 Modeling amino acid substitution

Due to the unavailability of the crystallized structure of TPO-R in RCSB Protein Data Bank, the structure of TPO-R was predicted by using the RaptorX server. The 3D analysis of wild type and mutant protein structures was performed by using UCSF Chimera (Figure 2-6).

**Figure2:**
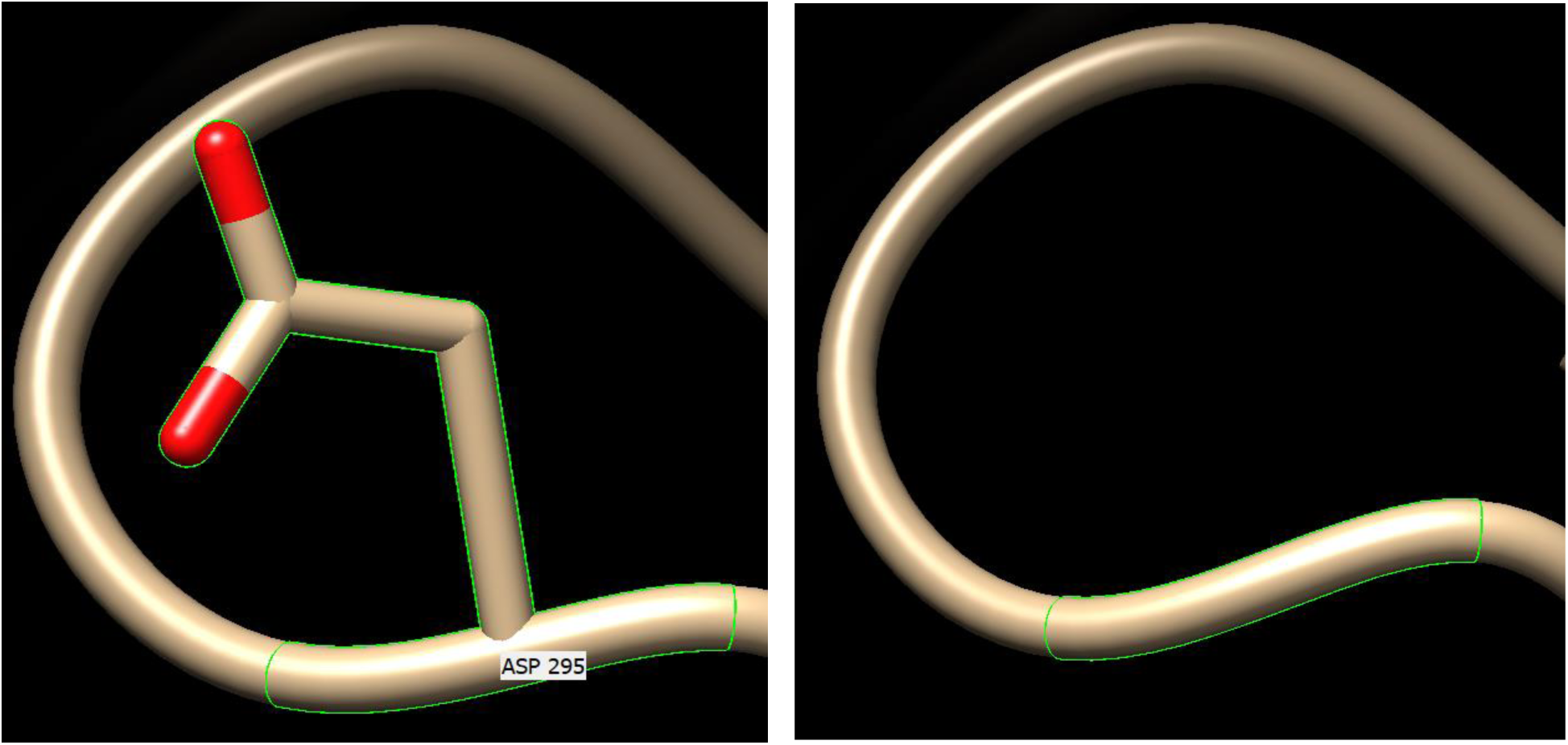
Showing 3D structure of D295G substitution using Chimera software. The wild-type (left) and the mutant residue (right). SNP ID: rs113696793, protein position 295 changed from Aspartic acid to glycine.

**Figure3:**
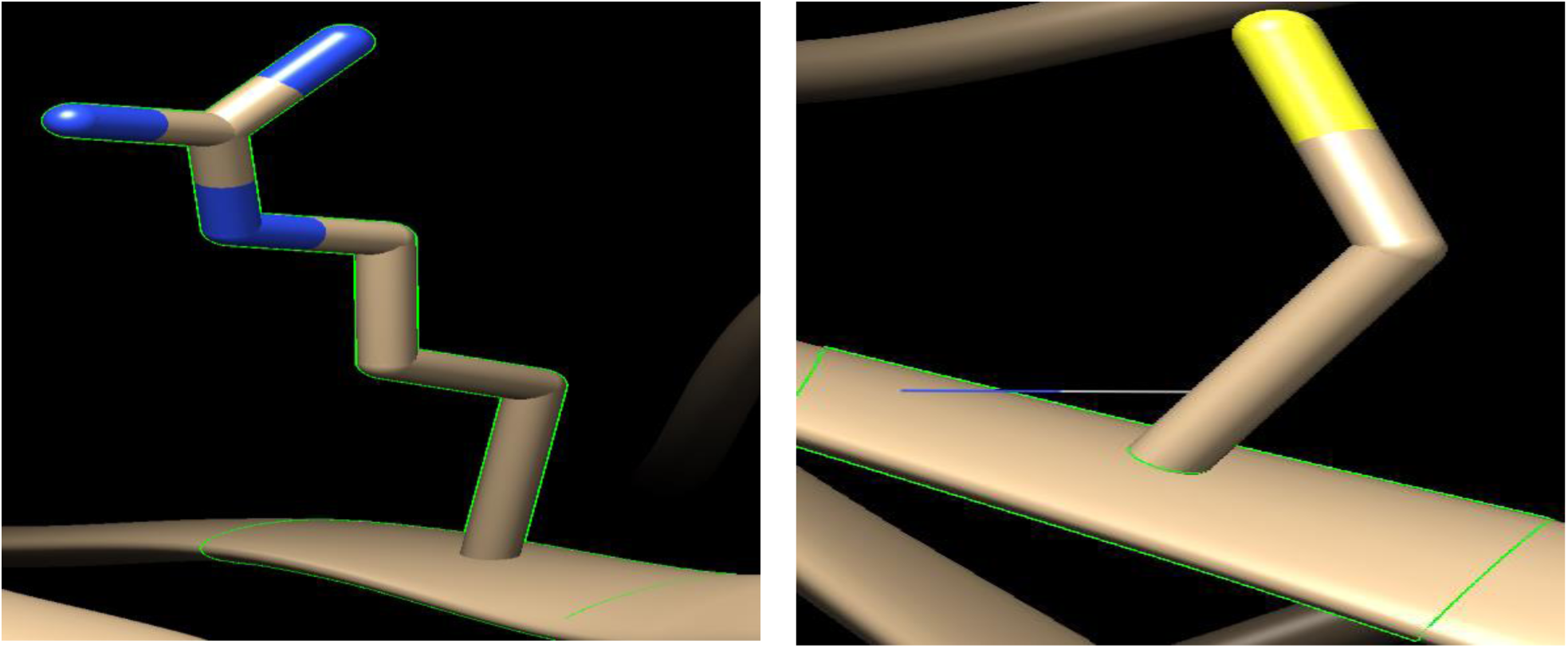
Showing 3D structure of R257C substitution using Chimera software. The wild-type (left) and the mutant residue (right). SNP ID: rs121913611, protein position 257 changed from Arginine to Cysteine.

**Figure4:**
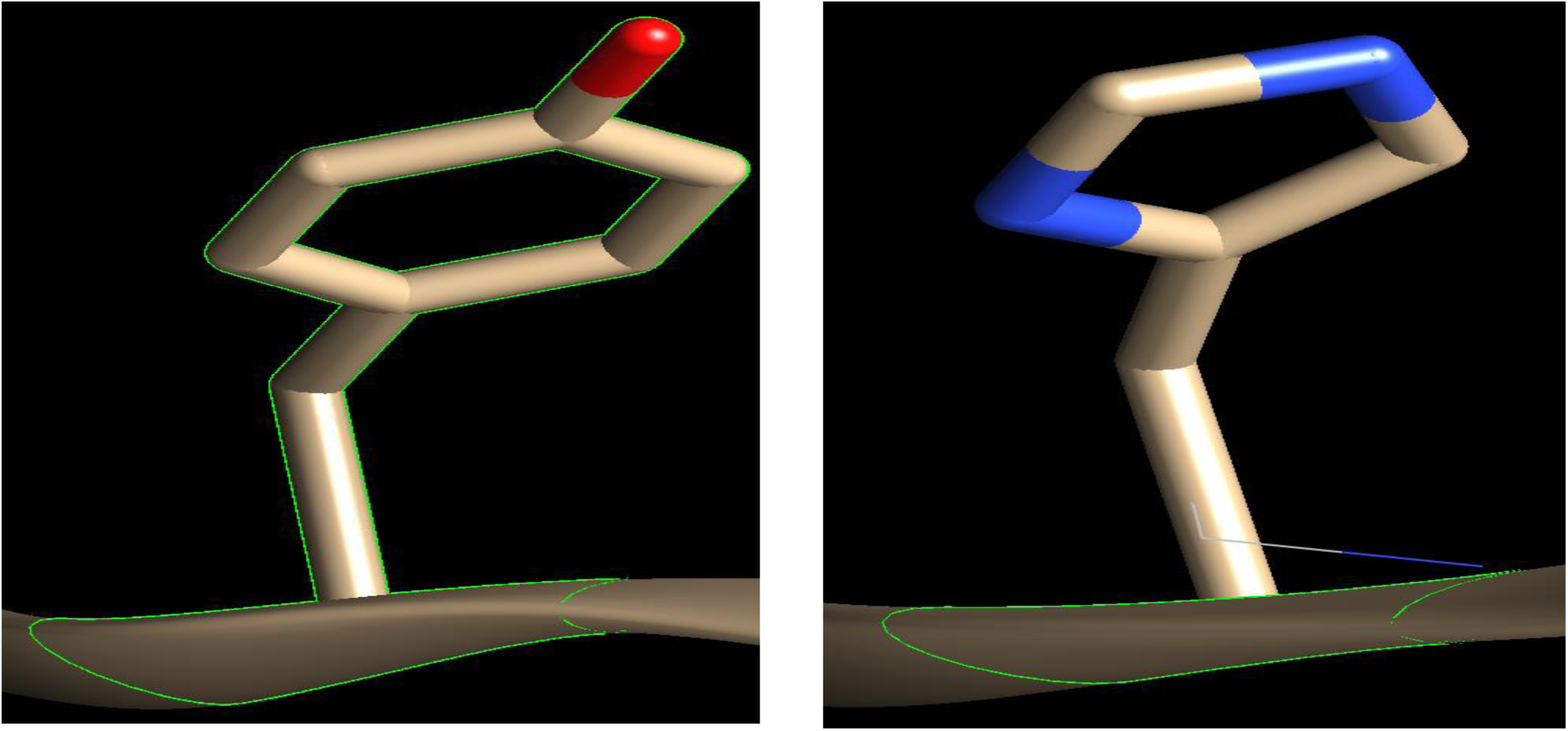
Showing 3D structure of Y252H substitution using Chimera software. The wild-type (left) and the mutant residue (right). SNP ID: rs141311765, protein position 252 changed from Tyrosine to Histidine.

**Figure5:**
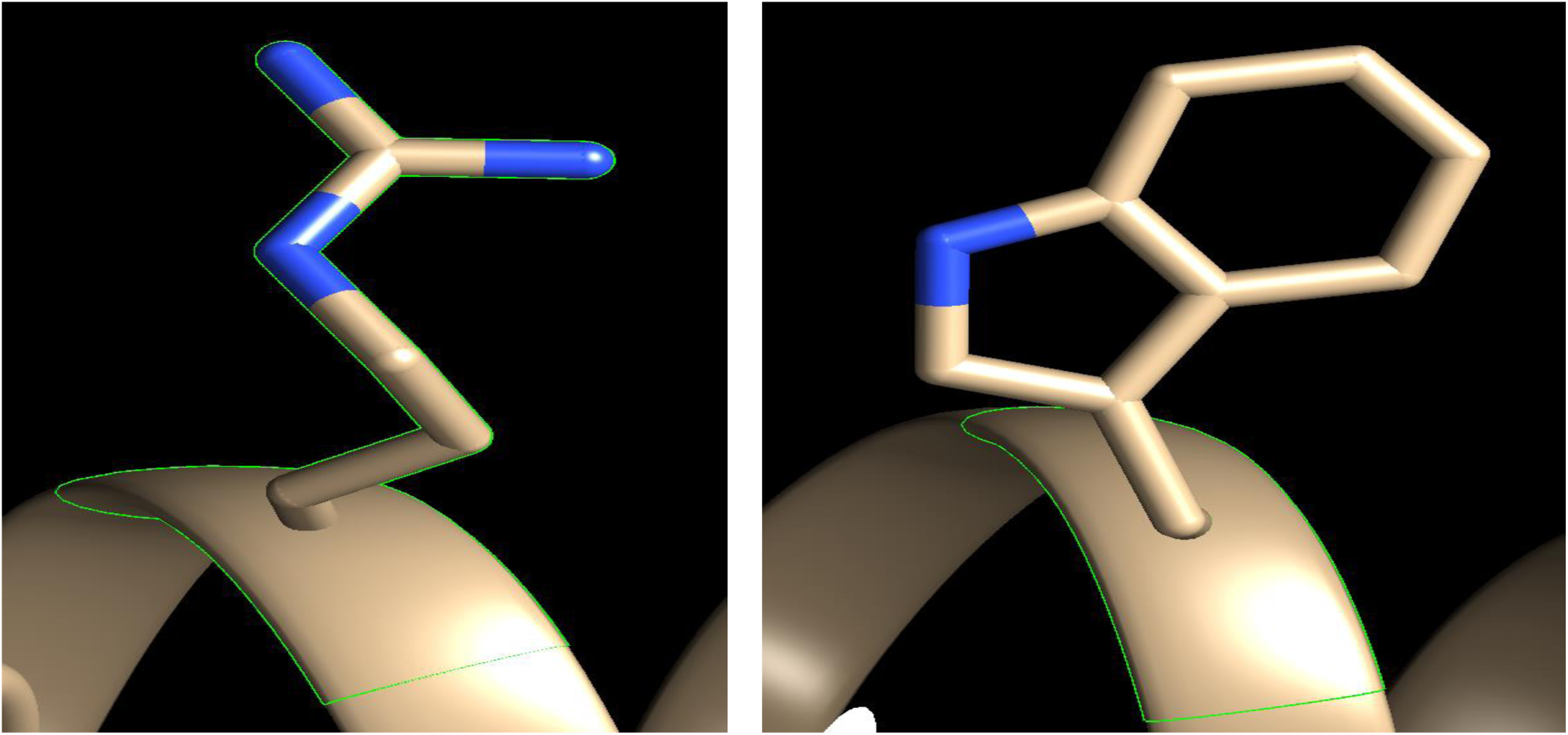
Showing 3D structure of R537W substitution using Chimera software. The wild-type (left) and the mutant residue (right). SNP ID: rs148784027, protein position 537changed from Arginine to Tryptophan.

**Figure6:**
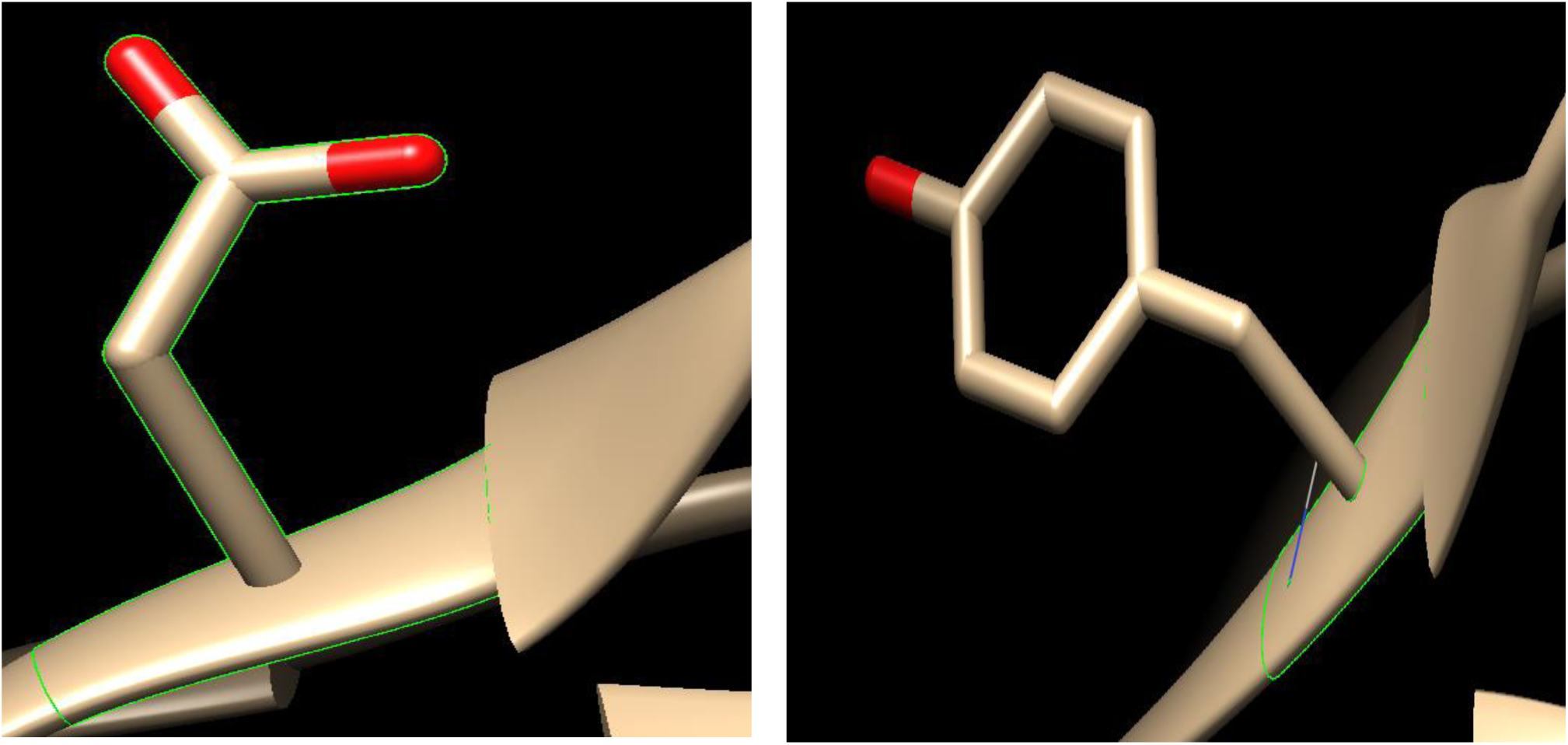
Showing 3D structure of D128Y substitution using Chimera software. The wild-type (left) and the mutant residue (right). SNP ID: rs201101813, protein position 128 changed from Aspartic acid to Tyrosine.

### 3.4. Protein-protein interaction network analysis

STRING analysis revealed that MPL has strong interactions with 10 proteins including, GRB2, IL3, JAK2, PIK3CA, STAT3, STAT5A and STAT5B (Fig. 7).

**Figure 7:**
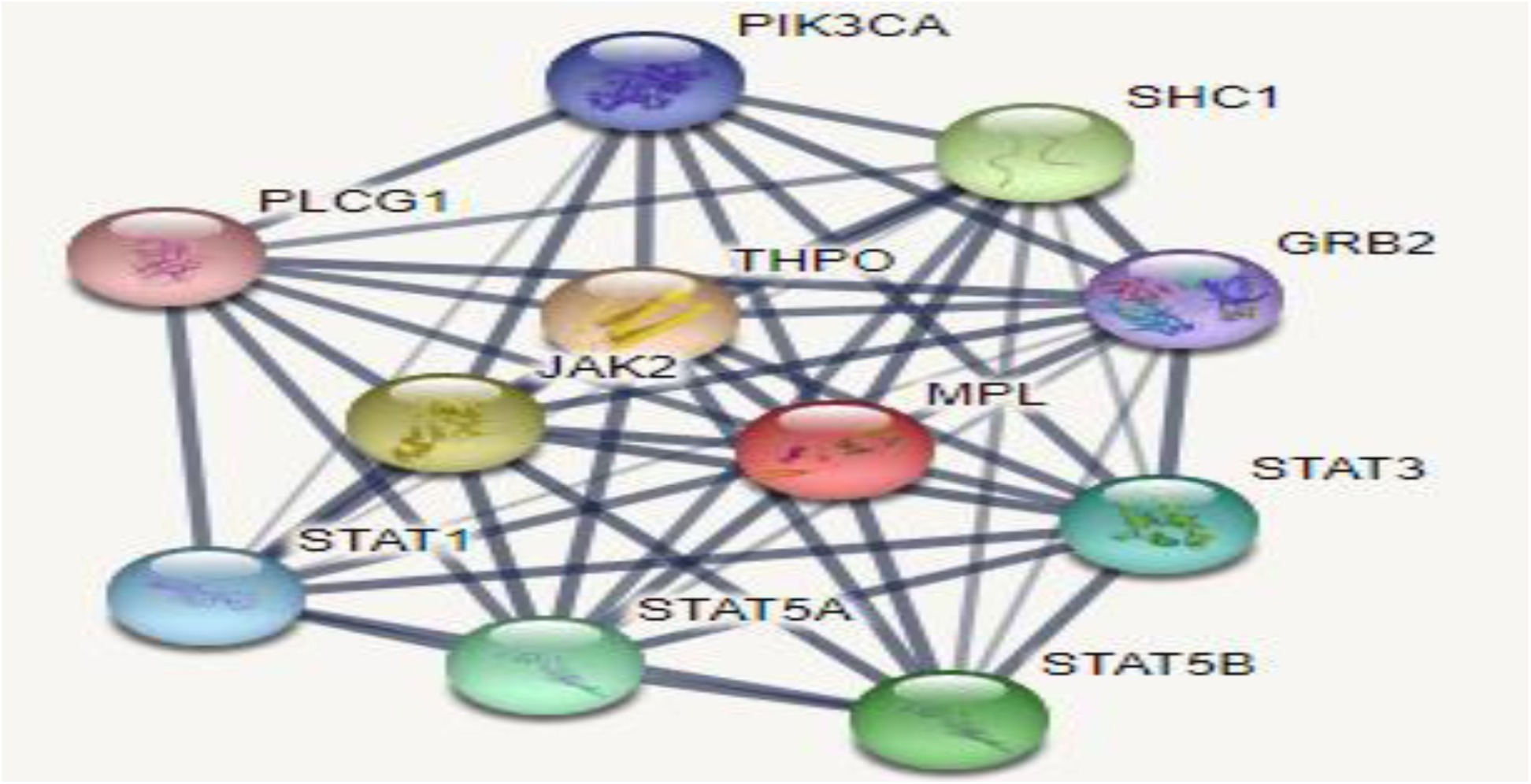
Protein–protein interaction network of TPO-R using STRING 9.0 server.

KEGG is a unique database source to simulate the functional networking of TPO-R in the form of molecular reaction and interaction networks, including metabolic pathways. KEGG showed the signaling transduction pathway for TPO-R in JAK-STAT, MAPK and PI3K-Akt signaling pathways (KEGG ID: <u>hsa04630</u>), (Figure. 8).

**Figure 8:**
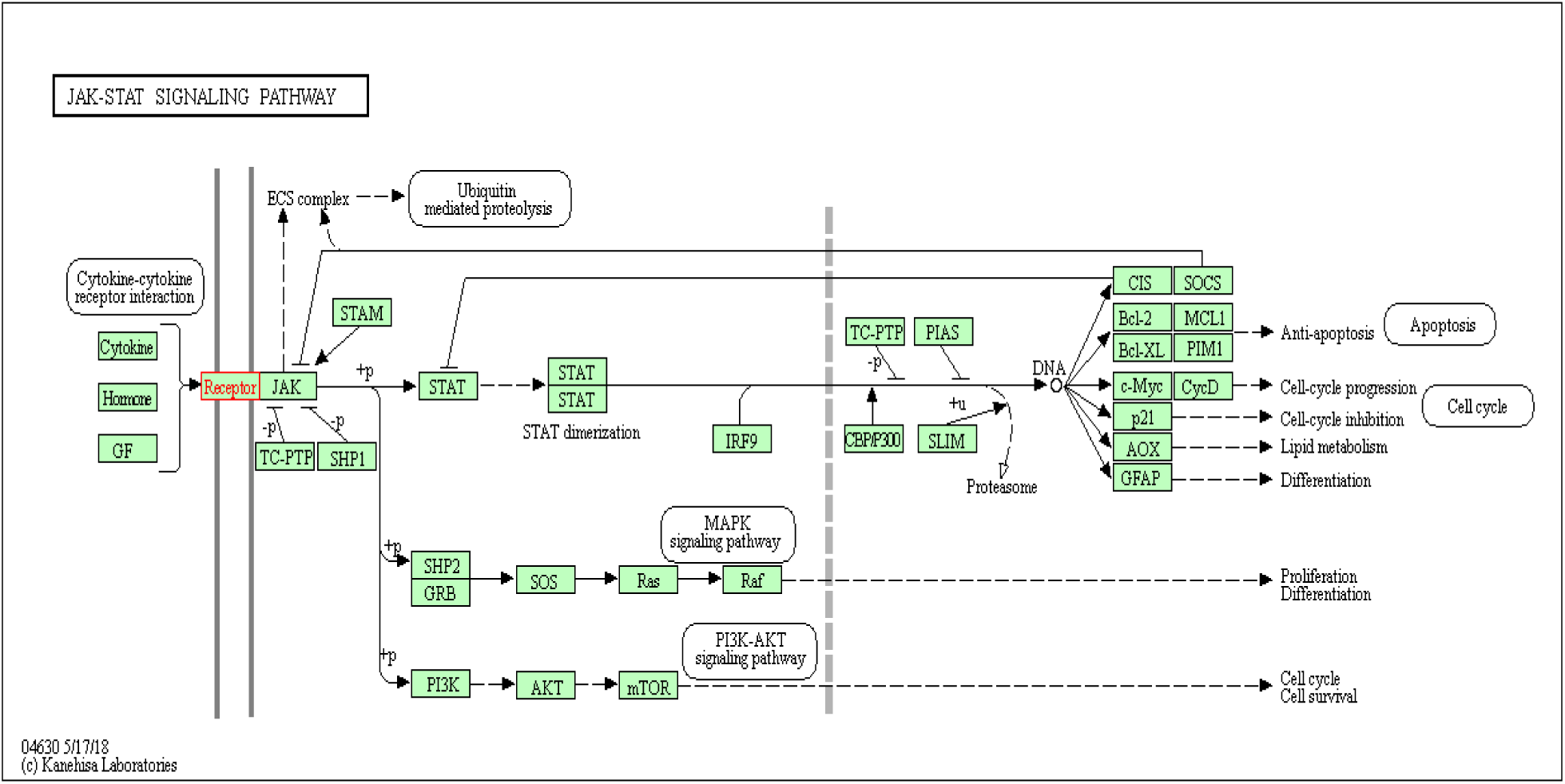
JAK-STAT pathway (KEGG ID: hsa04630)

## 4. Discussion

The current study undertakes a systematic in silico approach to screen the of functional mutation in human MPL gene for a better understanding of how do these mutations affect the protein function and structure and hence promote a disease.

On this in silico approach we select SIFT, Polyphen, PROVEAN, SNAP2, Condel, PhD-SNP, I-Mutant for the screening of functional mutation in MPL gene. By comparing the scores of all 6 methods, 5 nsSNPs with positions D295G, R257C, Y252H, R537W and D128Ywere found to be highly deleterious and decrease effective stability of the TPO-R.

The common physiochemical consequence of this nsSNPs were alteration in charge, size and hydrophobicity between wild-type and mutant residues which might lead to loss of interactions with other molecules and loss of hydrogen bonds in the core of the protein and as a result disturb correct folding.

The majority of post translational modifications effects this nsSNPs were altered disorderness in protein structure, gain or loss of catalytic residue, gain or loss of MoRF activity, alteration in phosphorylation sites and, gain of methylation, ubiquitination and glycosylation.

If the mutation predicted to altered disorderness in protein structure, the mutation will have considered likely to cause disease[35]. The five most frequent mutations (R→W, R→C, E→K, R→H, R→Q) collectively accounting for 44% of all deleterious disorder-to-order transitions [36]. Altered protein disorder was observed for substitution D295G, R257C, Y252H, R537W and D128Y.

Catalytic residues are defined as amino acid residues directly involved in the chemistry of catalysis contributing to substrate binding and protein stability [37]. Therefore, the gain of catalytic residues may change the rate of enzymatic activity and catalytic reaction. There is an evidence in the literature suggest that the loss and the gain of catalytic residues, may be actively involved in human-inherited disease [38]. Gain of catalytic residue was observed for substitutions R257C, Y252H and R537W.

Molecular recognition features (MoRFs) are short interaction prone segments intrinsically disordered regions (IDPs) in proteins sequences that fold upon binding to their interaction partners [39]. MoRFs are involved in protein-protein interactions, which serve as the initial step in molecular recognition and binding functions[40]. Intrinsically disordered proteins (IDPs) have been implicated in a number of human diseases, including cancer, diabetes and neurodegenerative [41-43]. Loss of MoRF binding activity was observed for substitution R537W, gain of MoRF binding activity was observed for substitution D128Y

Phosphorylation of proteins is an important regulatory mechanism as it acts as their molecular switch to perform various functions such as proliferation, apoptosis, regulation of metabolism, activating some proteins and deactivating others, and during signal transduction pathways [44]. Variation in the phosphorylation sites can change the enzymatic activity of a protein, disrupt the interactions between two or more proteins, changing the subcellular location of the phosphorylated protein [45]. There is an evidence in the literature that both gain and loss of a phosphorylation site may represent a molecular cause of disease for a number of inherited and somatic mutations [46, 47]. Loss of phosphorylation was observed for substitution R257C, gain of phosphorylation was observed for substitution D128Y.

Ubiquitylation is a post translational modifications which regulates several cellular mechanism such as protein degradation, cell cycle division, the immune response [48, 49]. Variation in the ubiquitination sites affects the degradation, alters cellular location of proteins, changes the protein activity, and alters protein interactions [50, 51]. The abnormality of ubiquitylation is linked to human pathologies varying from inflammatory neurodegenerative diseases to different forms of cancers [52,53]. Gain of ubiquitination was observed for substitution D295G making the region prone to degradation.

Protein methylation is post translational modifications mechanism plays role in the epigenetic control of gene expression, genome stability, signaling pathways and regulation of protein–protein interactions. [54,55]. Numerous studies reported that Methylated proteins, are involved in several human such as cancer [56,57] Gain of glycosylation was observed for substitution D295G.

Protein glycosylation is acknowledged as one of the major post-translational modifications, with significant effects on protein folding, conformation, distribution, stability and enzyme activity [58,59]. Alterations in glycosylation are common in lung and brain cancer and are thought to contribute to disease [60,61]. Gain of glycosylation was observed for substitution D128Y.

Here we present the results upon each residue and discuss the conformational variations.

### rs113696793

(D295G) this mutation resulted in a change of the aspartic acid to glycine at position 295. Glycines are very flexible and can disturb the required rigidity of the protein at this position. the aspartic acid residue charge was negative; the glycine residue charge is neutral. The Glycine residue is smaller and more hydrophobic than the wild-type residue. This residue is part of an interpret domain named “immunoglobulin-like fold” (IPR013783). Immunoglobulin(-like) domains are involved in a wide variety of functions which usually requires interaction of the intact domain with another protein/molecule. A mutation in such a domain could disturb this interaction.

As predicted by Mutpred, this substitution resulted in loss of sheet, loss of stability and loss of solvent accessibility but generate sites for ubiquitination and methylation. Based on conservation scores, D295is a highly conserved and exposed residue, this mutation thus making the TPO-R dysfunctional.

### rs121913611

(R257C) this mutation resulted in a change of arginine into a cysteine at position 257. The arginine residue charge was positive; the cysteine residue charge is neutral. The cysteine residue is smaller and more hydrophobic than the arginine residue. This residue is part of an interpret domains named “immunoglobulin-like fold” (IPR013783), “fibronectin type Iii” (IPR003961), and fibronectin type Iii Superfamily (IPR036116). The last two domains are annotated with the Gene-Ontology (GO) terms to indicate its function: protein binding (GO:0005515) and molecular function (GO:0003674). The effect of this variant is annotated as: Congenital amegakaryocytic thrombocytopenia (CAMT) [MIM:604498]. R257C results in gain of catalytic residue at R257 but loss of loop, loss of disorder and loss of phosphorylation of serine residue at 258^th^ as predicted by Mutpred. Based on conservation scores, R257is a highly conserved and exposed residue. This mutation is probably leading to the loss of activity of TPO-R.

### rs141311765

(Y252H) this mutation resulted in a change of the tyrosine to histidine at position 252. The histidine residue is smaller and less hydrophobic than the tyrosine residue, this might lead to loss of hydrophobic interactions in the core of the protein. This residue is part of an interpret domains named “Immunoglobulin-Like Fold” (IPR013783) “Fibronectin Type Iii” (IPR003961),and Fibronectin Type Iii Superfamily (IPR036116). The last two domains are annotated with the following Gene-Ontology (GO) terms to indicate its function: Protein Binding (GO:0005515)and Molecular Function (GO:0003674). Y252H mutation might disturb this function. This substitution also results in gain of protein disorder and gain of sheet but results in loss of stability and loss of loop. Moreover, this substitution also results in gain of catalytic residue at L254. Based on conservation scores, Y252 is a highly conserved and buried in the core of TPO-R, thus this mutation is probably disrupting structure of the TPO-R. A literature search revealed that Y252H has been shown to increase the thrombopoietin sensitive and associated with Essential thrombocythemia [62].

### rs148784027

(R537W) this substitution resulted in change of arginine (basic amino acid) to tryptophan (a non-polar aromatic amino acid). The arginine residue charge was positive; the tryptophan residue charge is neutral. The mutant residue is bigger and more hydrophobic this might. By loss of arginine, MoRF binding activity was found to be lost. It also leads to gain of catalytic activity at R537 residue but loss of disorder, loss of sheet and loss of solvent accessibility. Based on conservation scores, R537 highly conserved and exposed, this mutation is probably damaging to the TPO-R.

### rs201101813

(D128Y) this mutation involves the substitution of aspartic acid into a tyrosine at position 128. the aspartic acid residue charge was negative. the charge of the buried wild-type residue is lost by this mutation. The tyrosine residue is bigger and more hydrophobic than aspartic acid residue. This residue is located within an interpret domains named “immunoglobulin-like fold” (IPR013783), growth hormone/erythropoietin receptor, ligand binding (IPR015152) and fibronectin type Iii superfamily (IPR036116). These domains are important for binding of other molecules. The D128Y mutation might disturb the interaction between these domains and as such affect the function of the protein. The majority of physiochemical properties of this substitution were gain of phosphorylation at D128, gain of glycosylation at P133 and gain of MoRF binding activity. Based on conservation scores, D128 is a conserved and exposed residue. This mutation hence making the TPO-R nonfunctional.

The main signaling networks downstream of Tpo/ TPO-R include the JAK-STAT, MAPK and PI3K-Akt pathways. These pathways control essential cellular processes like differentiation, proliferation, cell survival and apoptosis with well-established roles in the initiation and progression of hematologic malignancies [63-65]. Interaction between the JAK-STAT pathway and PI3K-Akt signaling pathway has been reported in hematologic malignancies such as myeloid leukemia B-precursor ALL cells, leukemia and lymphoma [66,67].

## 5. Conclusions

Due to the importance of Tpo/TPO-R in survival, proliferation, and differentiation of hematopoietic cells, an extensive in silico analysis was performed to explore the potential effects of MPL genetic variants on functions and structure of TPO-R. Out of445 missense nsSNP, 5nsSNP were highly deleterious affect conserved positions and posttranslational modification sites of the TPO-R leading to loss or disturbance of internal and external interactions and ultimately loss of the function as well as the structure thereby, leading to diverse pathological conditions like low platelet levels, high serum TPO accompanies the thrombocytopenia. Four nsSNP rs113696793 (D295G) and rs121913611 (R257C) rs141311765 (Y252H) and rs201101813 (D128Y) lie in extracellular domain of TPO-R; therefore, these mutations may affect its binding to Tpo, subsequent probable disturb the Tpo/TPO-R downstream signaling pathways. One significant observation was the identification rs141311765 (Y252H), that has been shown to increase the thrombopoietin sensitive and associated with Essential thrombocythemia. The amino acid substitutions R537W residue is located within the cytoplasmic domain resulting in failure of signal transduction and JAKs binds JAKs and signal transduction. Furthermore, protein-protein interaction pathway has helped us to understand the roles of Tpo/TPO-R signaling pathways in differentiation, proliferation, cell survival and apoptosis of hematopoietic cells.

## References

1. Campbell PJ, Green AR. The myeloproliferative disorders. N Engl J Med. 2006;355:2452–66.

2. Hasselbalch HC, Bjorn RnME. MPNs as inflammatory diseases: the evidence, consequences, and perspectives. Mediators Inflamm. 2015;2015:e102476.

3. Vainchenker W, Kralovics R. Genetic basis and molecular pathophysiology of classical myeloproliferative neoplasms. Blood. 2017;129:667–79.

4. Cervantes F, Tassies D, Salgado C, et al. Acute transformation in nonleukemic chronic myeloproliferative disorders: Actuarial probability and main characteristics in a series of 218 patients. Acta Haematol. 1991;85:124–127.

5. Abdulkarim K, Girodon F, Johansson P, et al. AML transformation in 56 patients with Ph-MPD in two well defined populations. Eur J Haematol. 2009;82:106–111.

6. Baxter EJ, Scott LM, Campbell PJ, East C, Fourouclas N, Swanton S, et al. Acquired mutation of the tyrosine kinase *JAK2* in human myeloproliferative disorders. Lancet. 2005; 365:1054–1061.

7. Beer PA, Campbell PJ, Scott LM, Bench AJ, Erber WN, Bareford D, et al. MPL mutations in myeloproliferative disorders: analysis of the PT-1 cohort. Blood. 2008; 112:141–149.

8. Nangalia J, Massie CE, Baxter EJ, et al. Somatic CALR mutations in myeloproliferative neoplasms with nonmutated JAK2. N Engl J Med. 2013;369(25):2391–2405.

9. Vigon I, Mornon JP, Cocault L, Mitjavila MT, Tambourin P, Gisselbrecht S, Souyri M: Molecular cloning and characterization of MPL, the human homolog of the v-mpl oncogene: identification of a member of the hematopoietic growth factor receptor superfamily. Proc Natl Acad Sci U S A. 1992; 89: 5640–5644.

10. Souyri M, Vigon I, Penciolelli JF, Heard JM, Tambourin P, Wendling F: A putative truncated cytokine receptor gene transduced by the myeloproliferative leukemia virus immortalizes hematopoietic progenitors. Cell. 1990; 63: 1137–1147.

11. Mignotte V, Vigon I, de Crevecoeur EB, Romeo PH, Lemarchandel V, Chretien S: Structure and transcription of the human c-mpl gene (MPL). Genomics. 1994; 20: 5–12.

12. Varghese L.N., Defour J.P., Pecquet C., Constantinescu S.N. The thrombopoietin receptor: Structural basis of traffic and activation by ligand, mutations, agonists, and mutated calreticulin. Front. Endocrinol. (Lausanne) 2017; 8:59.

13. Forsberg EC, Prohaska SS, Katzman S, Heffner GC, Stuart JM, Weissman IL. Differential expression of novel potential regulators in hematopoietic stem cells. PLoS Genet. 2005; 1:e28.10.

14. Yoshihara H, Arai F, Hosokawa K, Hagiwara T, Takubo K, Nakamura Y, et al. Thrombopoietin/MPL signaling regulates hematopoietic stem cell quiescence and interaction with the osteoblastic niche. Cell Stem Cell. 2007; 1:685–97.10

15. Qian H, Buza-Vidas N, Hyland CD, Jensen CT, Antonchuk J, Mansson R, et al. Critical role of thrombopoietin in maintaining adult quiescent hematopoietic stem cells. Cell Stem Cell. 2007; 1:671–84.10.

16. Kaushansky K. Lineage-specific hematopoietic growth factors. N. Engl. J. Med. 2006; 354:2034–2045.

17. Kaushansky K. Thrombopoietin: accumulating evidence for an important biological effect on the hematopoietic stem cell. Ann NY Acad Sci 2003; 996:39–43.

18. Antonchuk J, Hyland CD, Hilton DJ, Alexander WS. Synergistic effects on erythropoiesis, thrombopoiesis, and stem cell competitiveness in mice deficient in thrombopoietin and steel factor receptors. Blood 2004; 104:1306–1313.

19. Ding J, Komatsu H, Wakita A, Kato-Uranishi M, Ito M, Satoh A, Tsuboi K, Nitta M, Miyazaki H, Iida S. et al. Familial essential thrombocythemia associated with a dominant-positive activating mutation of the c-MPL gene, which encodes for the receptor for thrombopoietin. Blood. 2004;103:4198–4200

20. Teofili L, Giona F, Torti L, Cenci T, Ricerca BM, Rumi C, Nunes V, Foa R, Leone G, Martini M. et al. Hereditary thrombocytosis caused by MPLSer505Asn is associated with a high thrombotic risk, splenomegaly and progression to bone marrow fibrosis. Haematologica. 2010; 95:65–70.

21. P.C. Ng SH. Predicting the effects of amino acid substitutions on protein function. Annu Rev Genomics Hum Genet. 2006;7 61–80.

22. Hoi Y and Chan AP. PROVEAN web server: a tool to predict the functional effect of amino acid substitutions and indels. Bioinformatics. 2015; 31(16): 2745–2747.

23. I.A. Adzhubei SS, L. Peshkin, V.E. Ramensky, A. Gerasimova, P. Bork, et al. A method and server for predicting damaging missense mutations. Nat Methods. 2010; 7: 248–9.

24. Hecht L, Wass J, Kelly L, Clevenger-Firley E, Dunn C. SNAP-Ed Steps to Health Inspires Third Graders to Eat Smart and Move More. Journal of Nutrition Education and Behavior. 2013; 45(6): 800–2.

25. Adzhubei IA, Schmidt S, Peshkin L, Ramensky VE, Gerasimova A, Bork P, et al. A method and server for predicting damaging missense mutations. Nat Methods. 2010;7(4):248–249.

26. Capriotti E CRaCR. Predicting the insurgence of human genetic diseases associated to single point protein mutations with support vector machines and evolutionary information. Bioinformatics. 2006; 22: 2729–34.

27. E. Capriotti PF, R. CasadioI-Mutant 2.0: predicting stability changes upon mutation from the protein sequence or structure. Nucleic Acids Res: 2005; 33: W306–W310.

28. Ashkenazy H, Erez E, Martz E, Pupko T, Ben-Tal N. ConSurf 2010: calculating evolutionary conservation in sequence and structure of proteins and nucleic acids. Nucleic Acids Res. 2010; 38: W529–33.

29. Li B, Krishnan VG, Mort ME, Xin F, Kamati KK, Cooper DN, et al. Automated inference of molecular mechanisms of disease from amino acid substitutions. Bioinformatics. 2009; 25(21): 2744–50.

30. Venselaar H, Te Beek TA, Kuipers RK, Hekkelman ML, Vriend G. Protein structure analysis of mutations causing inheritable diseases. An e-Science approach with life scientist friendly interfaces. BMC bioinformatics. 2010; 11: 548.

31. Wang, S, Li, W, Liu, S, Xu, J. RaptorX-Property: a web server for protein structure property prediction. Nucleic Acids Res. 2016;44:W430–W435.

32. Pettersen, EF, Goddard, TD, Huang, CC et al. UCSF Chimera —a visualization system for exploratory research and analysis. J Comput Chem. 2004;25: 1605–1612.

33. Mering V. STRING: known and predicted protein-protein associations, integrated and transferred across organisms. Nucleic Acids Res 2005; 33(Suppl. 1): D433–D7.

34. Kanehisa, Minoru and Susumu Goto. KEGG: Kyoto Encyclopedia of Genes and Genomes. Nucleic acids research. 1999; 27 (1): 29–34.

35. Vacic V., Markwick P. R. L., Oldfield C. J., Zhao X., Haynes C., Uversky V. N. and Iakoucheva L. M. Disease-associated mutations disrupt functionally important regions of intrinsic protein disorder. PLoS Comput. Biol. 2012; 8, e1002709.

36. Vacic V, Iakoucheva LM. Disease mutations in disordered regions-exception to the rule? Mol Biosyst. 2012; 8: 27–32.

37. Bartlett GJ, et al. Analysis of catalytic residues in enzyme active sites. J. Mol. Biol. 2002; 324:105–121.

38. Fuxiao Xin, Steven Myers, Yong Fuga Li, David N Cooper, Sean D Mooney, Predrag Radivojac. Structure-based kernels for the prediction of catalytic residues and their involvement in human inherited disease. Bioinformatics. 2010; 26, 1975–1982.

39. Cheng Y., LeGall T., Oldfield C.J., Dunker A.K., Uversky V.N. Abundance of intrinsic disorder in protein associated with cardiovascular disease. Biochemistry. 2006; 45:10448–10460.

40. Kotta-Loizou I Tsaousis GN, Hamodrakas SJ. Analysis of Molecular Recognition Features (MoRFs) in membrane proteins. Biochim Biophys Acta. 2013 Apr;1834(4):798–807.

41. Iakoucheva L.M., Brown C.J., Lawson J.D., Obradovic Z., Dunker A.K. Intrinsic disorder in cell-signaling and cancer-associated proteins. J. Mol. Biol. 2002; 323:573–584.

42. Uversky V.N. The triple power of d(3): Protein intrinsic disorder in degenerative diseases. Front. Biosci. (Landmark Ed.) 2014; 19:181–258.

43. Du Z., Uversky V.N. A comprehensive survey of the roles of highly disordered proteins in type 2 diabetes. Int. J. Mol. Sci. 2017; 18:10.

44. Nishi H, Fong JH, Chang C, Teichmann SA, Panchenko AR. Regulation of protein-protein binding by coupling between phosphorylation and intrinsic disorder: analysis of human protein complexes. Mol Biosyst. 2013; 9:1620–1626.

45. Michael J. Lee and Michael B. Yaffe. Protein Regulation in Signal Transduction. Cold Spring Harb Perspect Biol. 2016; 8(6): a005918.

46. Hanahan D, Weinberg RA. Hallmarks of cancer: the next generation. Cell. 2011; 144:646–674.

47. Harsha HC, Pandey A. Phosphoproteomics in cancer. Mol Oncol. 2010; 4:482–495.

48. Kerscher, O., Felberbaum, R., & Hochstrasser, M. Modification of Proteins by Ubiquitin and Ubiquitin-Like Proteins. Annual Review of Cell and Developmental Biology. 2006; 22(1), 159–180.

49. Elsasser S, Gali RR, Schwickart M, Larsen CN, Leggett DS, et al. Proteasome subunit Rpn1 binds ubiquitin-like protein domains. Nat. Cell Biol. 2002; 4:725–30.

50. Schnell JD, Hicke L (September 2003). “Non-traditional functions of ubiquitin and ubiquitin-binding proteins”. The Journal of Biological Chemistry. 278 (38): 35857–60.

51. Mukhopadhyay D, Riezman H. Proteasome-independent functions of ubiquitin in endocytosis and signaling. Science. 2007;315 (5809): 201–5.

52. Qiuyang Zheng, Timothy Huang, Lishan Zhang, Ying Zhou, Hong Luo, Huaxi Xu, and Xin Wang. Dysregulation of Ubiquitin-Proteasome System in Neurodegenerative Diseases. Front Aging Neurosci. 2016; 8: 303.

53. L. H. Gallo, J. Ko & D. J. Donoghue. The importance of regulatory ubiquitination in cancer and metastasis. Cell Cycle. 2017; 16:7, 634–648.

54. Phillips, T. The role of methylation in gene expression. Nature Education. 2008; 1(1):116J

55. Winter DL1, Abeygunawardena D1, Hart-Smith G1, Erce MA1, Wilkins MR1. Lysine methylation modulates the protein-protein interactions of yeast cytochrome C Cyc1p. Proteomics. 2015;15(13):2166–76..

56. Coralie Poulard, Laura Corbo, and Muriel Le Romancer. Protein arginine methylation/demethylation and cancer. Oncotarget. 2016; 7(41): 67532–67550.

57. Dong Han, Mengxi Huang, Ting Wang, Zhiping Li, Yanyan Chen, Chao Liu, Zengjie Lei, Xiaoyuan Chu. Lysine methylation of transcription factors in cancer. Cell Death and Disease 2019; 10:290.

58. Hanson, S. R. et al. The core trisaccharide of an N-linked glycoprotein intrinsically accelerates folding and enhances stability. Proc. Natl. Acad. Sci. U. S. A. 2009;106, 3131–3136.

59. Skropeta, D. The effect of individual N-glycans on enzyme activity. Bioorg. Med. Chem. 2009; 17, 2645–2653.

60. Hassan Lemjabbar-Alaoui, Andrew McKinney, Yi-Wei Yang,1 Vy M. Tran, and Joanna J. Phillips. Glycosylation alterations in lung and brain cancer. Adv Cancer Res. 2015;126:305–44.

61. Hakomori SI, & Cummings RD. Glycosylation effects on cancer development. Glycoconjugate Journal. 2012; 29(8-9), 565–566.

62. Michele P. Lambert, Jing Jiang, Vandana Batra, Chao Wu, and Wei Tong. A novel mutation in MPL (Y252H) results in increased thrombopoietin sensitivity in Essential Thrombocythaemia. Am J Hematol. 2012; 87(5): 532–534.

63. Yao Z, Cui Y, Watford WT, Bream JH, Yamaoka K, Hissong BD, Li D, Durum SK, Jiang Q, Bhandoola A, Hennighausen L, O’Shea JJ. Stat5a/b are essential for normal lymphoid development and differentiation. Proc Natl Acad Sci U S A. 2006;103:1000–1005.

64. Gustavo Loureiro, Daniella Bahia Kerbauy, Maria de Lourdes, L. F. Chauffaille, Maria Lucia M. Lee, Franciele B. M. Candido, Marçal C.A Silva, Eliza Y. S. Kimura, Fabio P. Guirao, Denise C. Rezende and Mihoko Yamamoto. Evaluation of MAPK and PI3K/AKT Signaling Pathways In Adult Acute Lymphoblastic Leukemia. Blood. 2010; 116:3240.

65. Osamu Yamada and Kiyotaka Kawauchi. The role of the JAK-STAT pathway and related signal cascades in telomerase activation during the development of hematologic malignancies. JAKSTAT. 2013; 2(4): e25256.

66. Camille Malouf and Katrin Ottersbach. Molecular processes involved in B cell acute lymphoblastic leukaemia. Cell Mol Life Sci. 2018; 75(3): 417–446.

67. Chen E1, Staudt LM, Green AR. Janus kinase deregulation in leukemia and lymphoma. Immunity. 2012;36(4):529–41.

